# The electrophysiological underpinnings of variation in verbal working memory capacity

**DOI:** 10.1101/2020.05.02.073825

**Authors:** Yuri G. Pavlov, Boris Kotchoubey

## Abstract

Working memory (WM) consists of short-term storage and executive components. We studied cortical oscillatory correlates of these two components in a large sample of 156 participants to assess separately the contribution of them to individual differences in WM. The participants were presented with WM tasks of above-average complexity. Some of the tasks required only storage in WM, others required storage and mental manipulations. Our data indicate a close relationship between frontal midline theta, central beta activity and the executive components of WM. The oscillatory counterparts of the executive components were associated with individual differences in verbal WM performance. In contrast, alpha activity was not related to the individual differences. The results demonstrate that executive components of WM, rather than short-term storage capacity, play the decisive role in individual WM capacity limits.

## Introduction

Working memory (WM) – our ability to maintain and manipulate information in a short term – is not a uniform construct. Although several theoretical models compete for the best explanation ^1^, most of them converge on the idea that WM entails (1) short-term storage for memory content (which can be regarded as a separate processing unit or as activated parts of long-term memory), and (2) executive components which are responsible for attentional control, protection from interference, active processing and reorganization, or manipulation of information in the short-term storage ^2,3^. As regards the former, it should be distinguished from the ultrashort-term storage in iconic or echoic memory. As regards the latter, these executive components are what makes WM “working” and differentiates the WM in the proper sense from mere short-term memory. Most WM tasks in real life require both short-term storage and executive components. For example, keeping a phone number in mind until it is dialed requires transformation of verbal information into a sequence of button presses. The task also involves switching attention between the current number to dial and pressing buttons on the phone panel. A more complex example is the conversion of a shopping list into the optimal path while shopping. This task, in addition to the translation of the list into a sequence of spatial locations, also involves constant updating of information in WM.

Available electrophysiological data indicate that manipulation of information in WM is strongly related to theta oscillations (4-8 Hz) in the medial prefrontal and anterior cingulate cortex for manipulation of information in WM ^4–12^. Therefore, theta is thought to reflect engagement of the executive components of WM ^13^. In turn, posterior alpha activity (8-13 Hz) has been associated with short-term storage, and may directly underlie maintenance of information in WM ^14,15^. Alpha activity responds to increasing load reaching an asymptote at the levels of individual’s WM capacity ^16^. Thus, posterior alpha activity can be seen as a reflection of the short-term storage component of WM.

The oscillatory mechanisms supporting individual WM capacity limits remain largely understudied. To study individual differences, large samples are essential. However, most available studies built their conclusions on samples of less than 30 individuals ^16–25^. Better-powered studies (∼35 participants per group) yielded mixed findings ^11,26–28^. For instance, after a median split, only the high-performance group showed an increase of theta with load ^11,28^, suggesting that theta activity and the related executive functions are important for successful maintenance of WM. But another study did not find this relationship ^27^. The relationship between alpha and beta activity and individual WM capacity is even less consistent.

The components of WM may contribute to individual differences in a different scale ^29,30^. Here, we sought to provide solid data to set apart the contribution of oscillatory counterparts of short-term storage and executive components of WM to individual differences. To reach this goal, we analyzed EEG data from a large number of participants (N=156) in WM tasks of above average complexity, which required either only retention of letter strings (Retention task) or retention and manipulation with the letters (mental recombination to the alphabetical order, Manipulation task) (see Figure 1 for the experimental design). Increasing the complexity of the tasks allowed us to increase the variance, to avoid ceiling effect in performance and to better distinguish between low and high performers. The task with mental manipulations allowed us to assess the contribution of different WM components. Although even the simple retention of information involves executive components of WM, the executive load of retention is tangibly weaker than the load of mental manipulations with the same content. The manipulation task possesses all the properties that the retention task does but additionally, it involves executive components in a much larger extent. Thus, the differential score between the spectral power in the tasks allowed us to derive a neural index of executive components of WM – executive index. We hypothesized that (i) the power of alpha oscillations would be related to the capacity of the short-term storage component of WM, (ii) the power of frontal midline theta would be related to the efficiency of the executive components of WM, and (iii) the latter mechanism, but not the former one, would be related to the overall memory performance.

**Figure 1.**
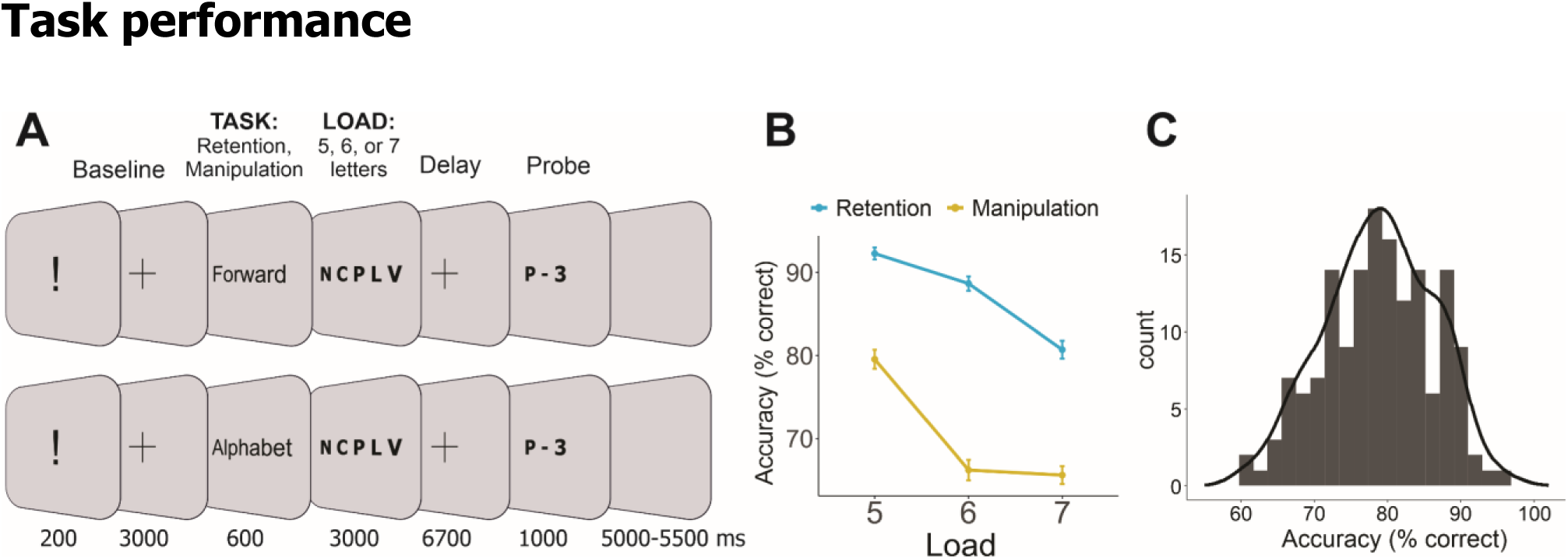
(A) The experimental paradigm. (B) Average performance in 6 conditions. Error bars are standard errors of the mean. (C) Distribution of overall performance stacked into 25 bins.

## Results

### Task performance

The accuracy was worse in the manipulation task and monotonically decreased with increasing WM load (main effect of Task: F(1, 155) = 431.8, η^2^ = 0.736, p < 0.001; Load: F(2, 307) = 138.8, η^2^ = 0.472, p < 0.001). The significant Task x Load interaction (F(2, 305) = 22.1, η^2^ = 0.125, p < 0.001) resulted from the fact that load increment from 6 to 7 letters yielded a strong drop of accuracy in the retention task but no effect in the manipulation task (see Figure 1B). All other pairwise differences were significant except the non-significant difference in performance between Retention of 7 and Manipulation of 5 letters.

### Time-frequency analysis

We employed the linear mixed effects models (LME) to explore the relationship between behavioral performance (quantified as average accuracy across all conditions) and the spectral power in the theta frequency band. Although we used LME with a continuous Performance variable for statistical inference, for the ease of data interpretation and illustration purposes, the whole sample was median-split into the high performance (N = 78) and low performance (N = 78) groups on the basis of their average accuracy across all conditions.

As can be seen in Figure 2, the high performance group exhibited a stronger theta increase in the Manipulation task than in the Retention task. This Task by Performance interaction (p = 0.004) resulted from the association between the relative theta power and behavioral performance being positive in the Manipulation task but negative in the Retention task (see Figure 2 and Supplementary Table S1 for full statistical output).

**Figure 2.**
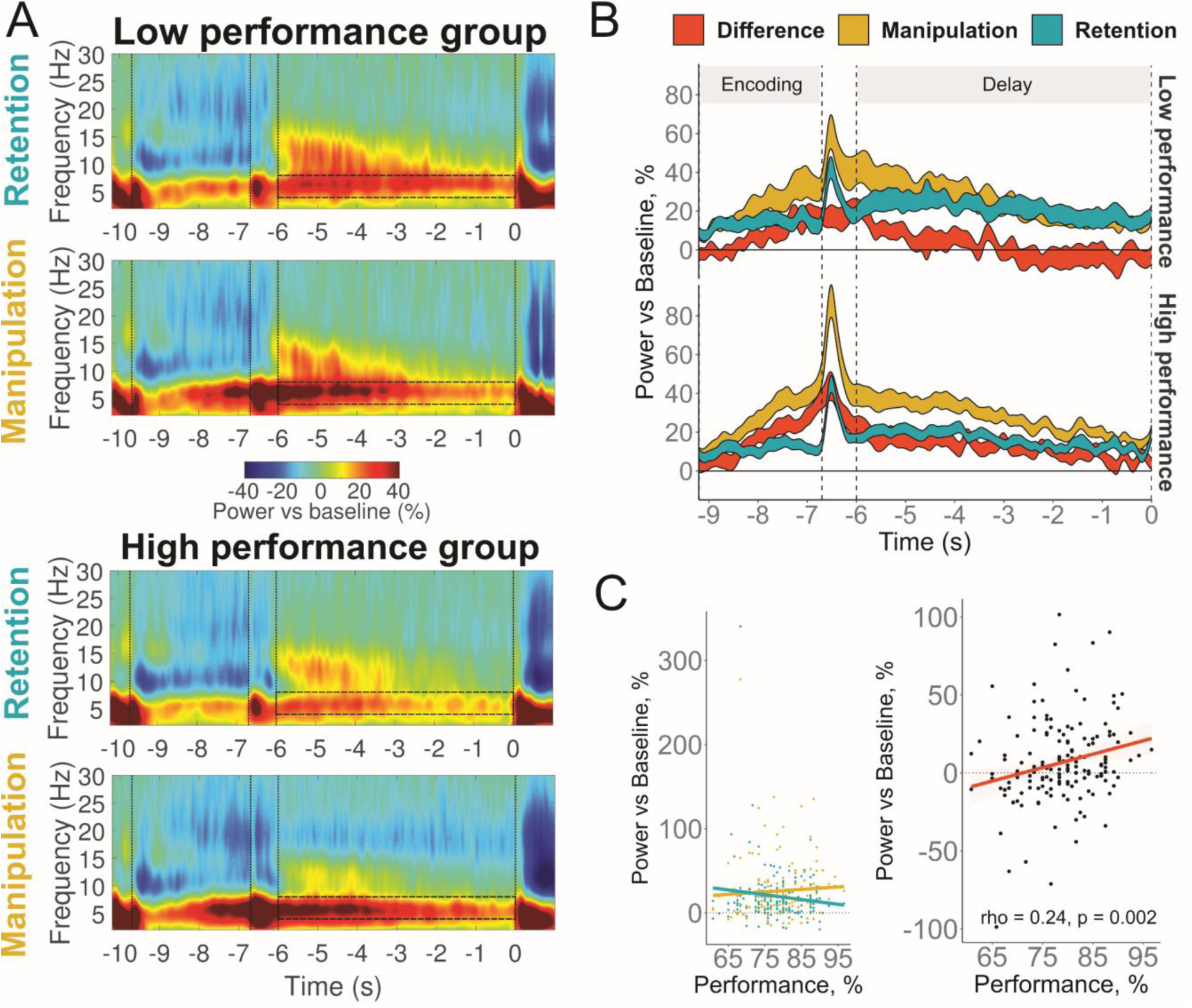
Relationship between theta power at Fz and WM performance. (A) Time-frequency maps in low (top panels) and high (bottom panels) performance groups in retention and manipulation tasks. Semi-transparent boxes mark time-frequency windows of interest (4-8 Hz, last 6 s of delay). The first vertical bar marks the onset of encoding; the second one marks the onset of delay; and the third one marks the beginning of the time-frequency window of interest; and the fourth one marks the onset of retrieval. (B) The time dynamics of theta activity in Retention, Manipulation tasks and their difference: the result of subtraction of theta relative power in the retention task from the power in the manipulation task (executive index). The width of the line represents 2 standard errors of the mean. (C) Left panel: correlation of WM performance in the manipulation and retention tasks. Right panel: correlation of WM performance and the executive index. Performance is the average percent accuracy in all conditions.

Then, we correlated WM performance with the difference between theta in Manipulation and Retention tasks to test the hypothesis on the role of executive components of WM in individual differences. The larger was the difference between baseline normalized theta power in Manipulation and Retention conditions, the better the individual WM performance (Spearman’s rho = 0.24, p = 0.002) (see Figure 2C and Supplementary Figure S5 for full correlation matrix).

No significant main effect (p = .359) or interactions with Performance (Task x Performance: p = .454) were found in the alpha frequency band (see Figure 3 and Supplementary Table S2).

**Figure 3.**
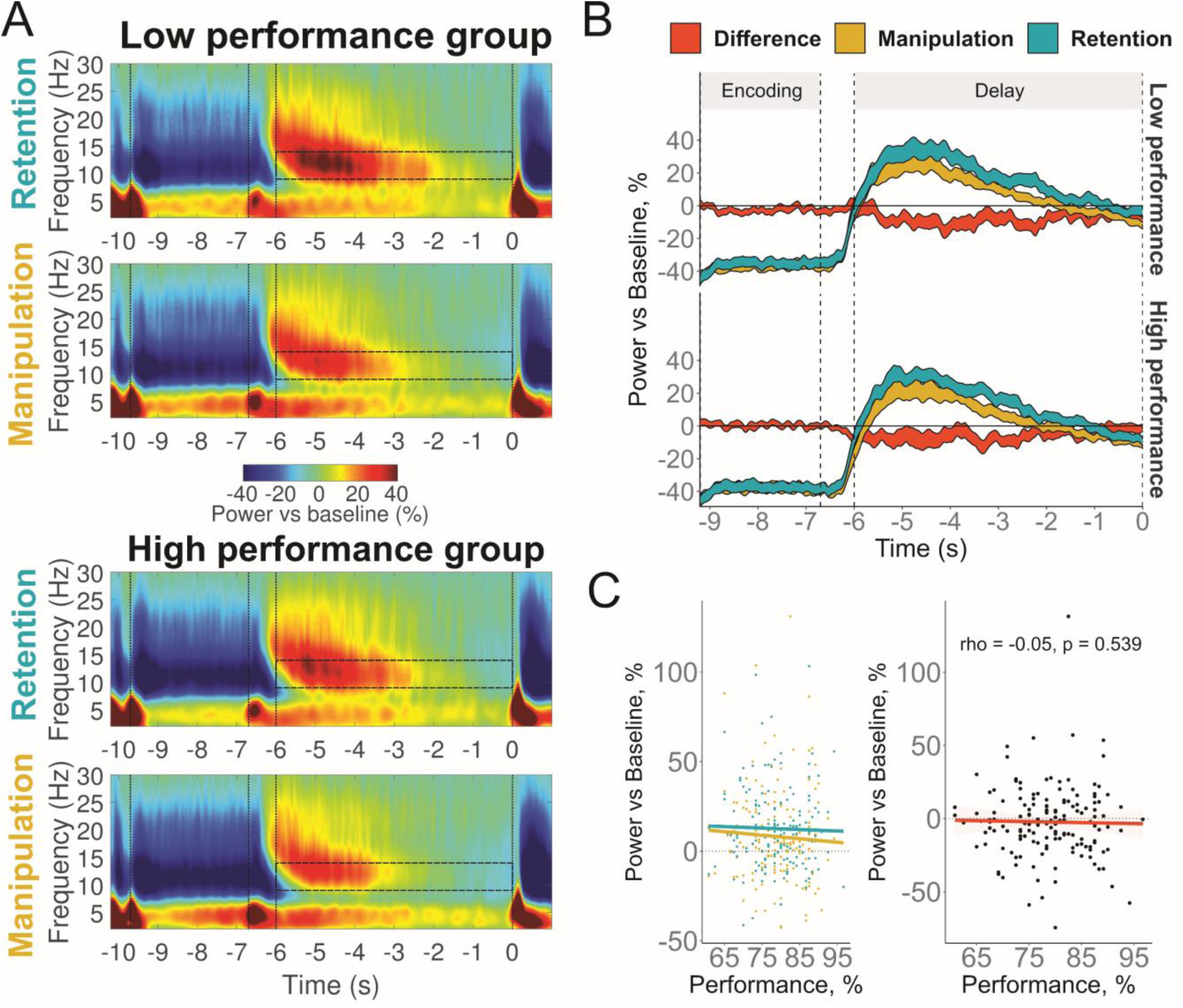
Relationship between alpha power over posterior ROI (P3, T5, O1, P4, T6, O2) and WM performance. (A) Time-frequency maps in low (top panels) and high (bottom panels) performance groups in retention and manipulation tasks. Semi-transparent boxes mark time-frequency windows of interest (9-14 Hz, last 6 s of delay). The first vertical bar marks the onset of encoding; the second one marks the onset of delay; and the third one marks the beginning of the time-frequency window of interest; and the fourth one marks the onset of retrieval. (B) The time dynamics of theta activity in Retention, Manipulation tasks and their difference: the result of subtraction of theta relative power in the retention task from the power in the manipulation task (executive index). The width of the line represents 2 standard errors of the mean. (C) Left panel: correlation of WM performance in the manipulation and retention tasks. Right panel: correlation of WM performance and the executive index. Performance is the average percent accuracy in all conditions.

Beta power negatively related to WM performance in the Manipulation task but not in the Retention task (Task by Performance interaction: p<0.001; see Figure 4 and Supplementary Table S3). Like it has been done with the theta activity, a subtraction of Retention beta from Manipulation beta was taken as an index of executive WM components. The correlation between this index and WM performance had similar magnitude as yielded in the analysis of theta (rho = −0.26, p = 0.001), but with the opposite sign (see Figure 4). Thus, better performance was related to lower beta activity.

**Figure 4.**
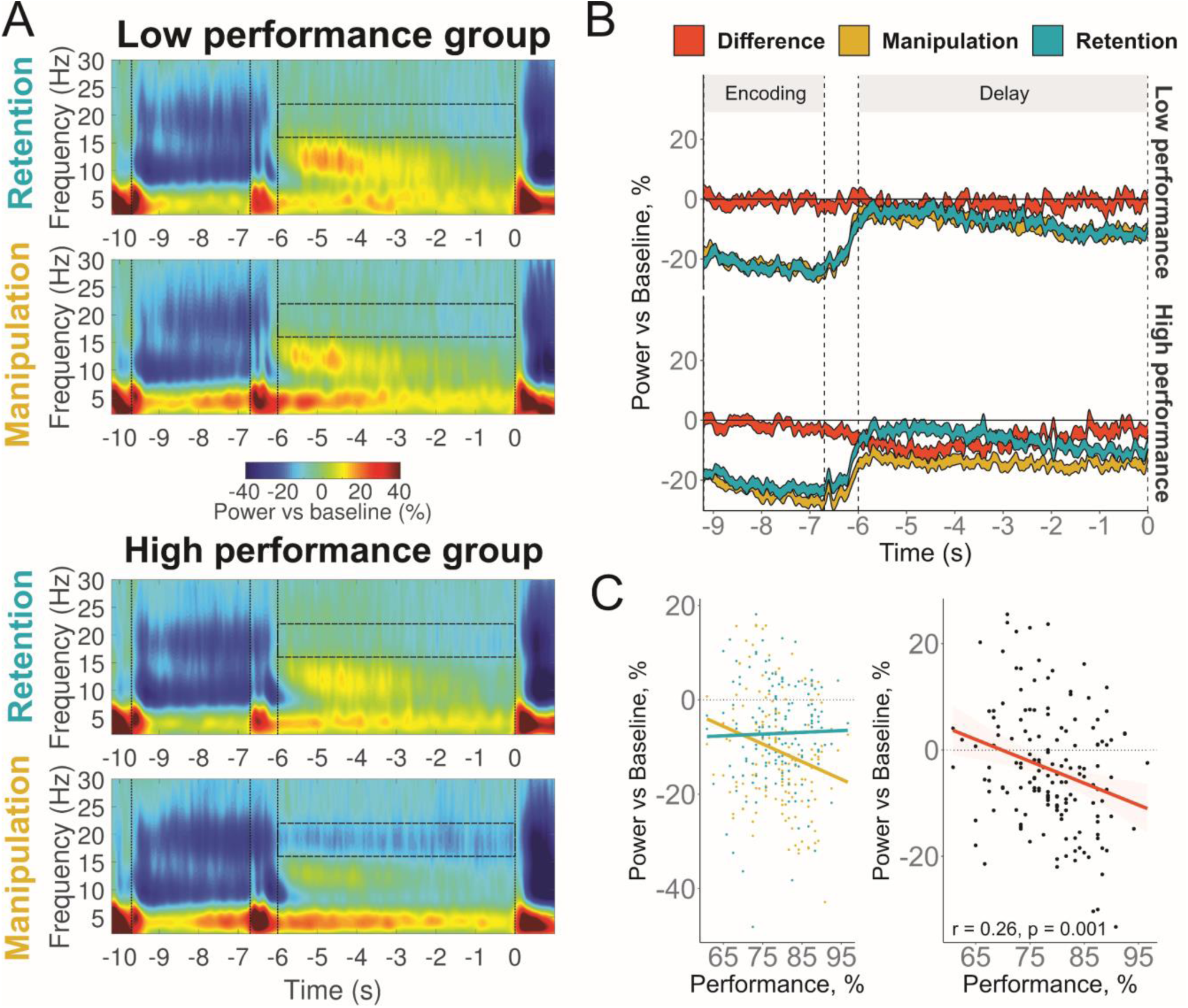
Relationship between beta power over central ROI (Cz, C4, C3) and WM performance. (A) Time-frequency maps in low (top panels) and high (bottom panels) performance groups in retention and manipulation tasks. Semi-transparent boxes mark time-frequency windows of interest (16-22 Hz, last 6 s of delay). The first vertical bar marks the onset of encoding; the second one marks the onset of delay; and the third one marks the beginning of the time-frequency window of interest; and the fourth one marks the onset of retrieval. (B) The time dynamics of theta activity in Retention, Manipulation tasks and their difference: the result of subtraction of theta relative power in the retention task from the power in the manipulation task (executive index). The width of the line represents 2 standard errors of the mean. (C) Left panel: correlation of WM performance in the manipulation and retention tasks. Right panel: correlation of WM performance and the executive index. Performance is the average percent accuracy in all conditions.

Because the correlations of beta and theta executive indices with performance are similar, we tested whether the two represent the same underlying mechanism. The correlation between the constructs was in an expected direction but not statistically significant (rho = −0.12, p = 0.124). Another way to test independence of the executive indices is to use Bayesian linear regression. Employing this approach, we, first, compared all three possible models – only difference score beta, only difference score theta, and a combination of both difference scores as IVs – with the intercept only model. Second, we compared the one predictor models with the two predictors model. The model preferred by Bayes factor is the two predictors model (BF = 135.48), and the model is 14.61 times better than only theta (BF = 9.27), and 3.85 times better than only beta model (BF = 35.23). The fact that the highest Bayes factor was yielded by the two predictors model suggests that their contribution is largely independent. Thus, theta and beta may play independent roles in individual differences in WM.

Then to check the reliability of the above effects we used a residual change score instead of difference as difference scores. Using a residual change scores for theta (rho = 0.24, p = 0.003) and beta (rho = −0.28, p < 0.001) successfully replicated the results obtained with the difference scores. The correlation between alpha and performance remained non-significant (rho = −0.06, p = 0.49).

## Discussion

We found that two EEG indices of the executive components of WM (calculated as the difference between the relative spectral power in the retention and the manipulation conditions or, alternatively, as residual change scores) significantly correlated with WM performance in a large sample of healthy individuals. These findings confirm our hypothesis (iii) stating that oscillatory associates of the executive WM component, but not of the storage component, are related to WM performance. This is congruent with previous behavioral research indicating that individual differences in executive functions were better predictors of WM capacity than differences in short-term storage ^30–32^.

The positive correlation between WM performance and the theta executive index, (i.e., more theta, better performance) also confirms our hypothesis (ii). Although in STM tasks the demand on executive functions is considerably lower than in genuine WM tasks, it is not completely lacking. For example, when WM load is high, executive components of WM are needed to counteract interference and to suppress irrelevant information. Few studies demonstrated a correlation between theta activity and performance in verbal (Zakrzewska & Brzezicka (2014): N=69, r=0.32, Kwon et al. (2015): N=13, r=0.76) and visual (Kawasaki & Yamaguchi (2013): N=14, r=0.51; Maurer et al. (2015): N=24, r=-0.41) short-term memory tasks that require only retention of, but no manipulation with, stimulus material. In contrast, we found no correlation between theta activity in the Retention task and behavioral performance (rho = −0.01, p = 0.88). Due to the known effects of saturation of brain activity measures with reaching higher levels of load ^16,33–35^, we hypothesized that the role of theta in retention tasks may be limited to only lower levels of load. However, even at the lowest level of 5 letters the correlation between theta during retention and the accuracy was not significant (rho = −0.03, p = 0.67). Thus, theta increase during delay in simple tasks was not predictive to performance in genuine WM tasks.

Contrary to theta, the beta index of the executive control negatively correlated with WM performance (i.e., more beta, worse performance). This finding was not predicted by any hypothesis because the literature concerning this relationship is too scarce. Probably the only comparable study is a spatial WM MEG experiment ^36,37^. Both publications reported a negative correlation between beta (15-20 Hz) and behavioral performance. Although the two reports shared the same sample, the localization of the correlation was different: WM performance correlated with beta activity in the right superior parietal lobule in one paper, and in the left dorsolateral prefrontal cortex and the bilateral superior temporal gyrus in the other. Moreover, beta activity in the two reports correlated with two different indexes of WM performance. These inconsistencies cast doubt on the reliability of the results.

Beta desynchronization may reflect switching internal attention between the reorganized set of letters and the initial set during the manipulations. Even when attention shifts from one object to another occur without eye movements, the shifts activate the same cortical areas as real eye movements ^38^. A good example is the activation of frontal eye fields (FEF) found in a spatial manipulation task ^39^. If the beta activity observed in our study is related to activation of motor control networks, then why was it related to individual differences in WM? Repetitive saccade eye movements during delay were shown to increase episodic memory performance ^40,41^. The movements executed just before blocks of an attentional control task (Flanker) also improved performance in the task ^42^. The attentional enhancement by preactivation of fronto-parietal network nodes such as FEF was hypothesized to produce the above mentioned effects ^42^. Consistent with this idea, TMS delivered to FEF improved detection of targets in a visuospatial attention task ^38^, whereas a suppression of the same region by TMS decreased inhibitory control ^43^. It is admittedly a speculation at this point, but more efficient use of FEF to execute mental manipulations would potentially facilitate WM performance in our study.

Our hypothesis (i) predicted a significant link between alpha activity and the short-term storage capacity. This hypothesis was not confirmed. Although this link can be expected on the basis of theoretical considerations (as mentioned above in the Introduction), empirical data are not very consistent. Correlations between WM capacity and posterior alpha can be strongly negative (Kawasaki & Yamaguchi,2013: visual WM, N = 14, r = −0.66) or strongly positive (Kwon et al., 2015: verbal WM, N = 13, r = 0.75). Another study found no correlation between WM performance and relative alpha power in the Sternberg task (Sghirripa et al., 2019; N=24). Better WM performance was associated, in some studies, with stronger alpha enhancement in more difficult tasks than in the easy tasks (Hu et al., 2019: N = 20, r = 0.55; N = 23, r = 0.59), in other studies, however, with stronger alpha suppression in more difficult tasks (Erickson et al., 2019: N = 60, r = 0.45; Fukuda et al., 2015: N = 28, r = 0.48). In a yet another study the positive correlation between the WM capacity and alpha modulation from lower to the higher load did not reach significance (Roux et al., 2012: N=25, r=0.39). We calculated the alpha modulation index (within-subject linear regression coefficients of alpha power by Load), as used in the verbal WM study by Hu et al. ^19^. The test attained significance neither for the overall task performance (rho = 0.04, p = 0.64), nor for the retention task only (rho = 0.03, p = 0.72).

Therefore, not all correlations obtained in the previous studies were replicated in a larger sample. One might suggest that our negative result is due to using only high-load conditions, and that stronger correlations would be obtained if we compared alpha between high-load and low-load conditions. Nevertheless, the individuals with higher WM capacity did not show signs of larger short-term storage that would have been reflected in the modulation of alpha at the high levels of load. We suggest that instead it is the executive control as reflected in the theta activity that controls attention and minimizes the interference, thus determining an individual’s WM capacity.

The interference hypothesis of WM capacity assumes that WM is affected not only by external distractors but also by mutual interference between the items ^44–46^. Thus, the detrimental effect on memory precision is a function of the number of items stored in WM and the presence of the concurrent task. When WM load is high and/or manipulations are required, the executive component of WM is needed to cope with interference. Perhaps, the resources that can be deployed to counteract interference are better represented in the higher performing individuals.

There is another possible mechanism to explain how executive processes may cope with increasing WM demands. Camos & Barrouillet ^47^ suggested that attention is involved in rapid refreshment of memory traces, thus preventing temporal decay of the WM content. The non-refreshed information fades out and in the worst case cannot be recovered. In extension of this model, beta and theta activity reflect the ability to maintain information in the active state by switching the focus of attention between items, thus defining the individual capacity limit.

Above we interpreted changes in posterior alpha rhythm as a reflection of the demand on the short-term storage component of WM. Another interpretation regards alpha as a mechanism of filtering information by gating only relevant sensory stimuli ^48,49^. If we accept this view, then either higher WM capacity individuals are no more efficient in the filtering out task-irrelevant information than lower WM capacity individuals, or the alpha filtering mechanism works only under moderate levels of memory load.

To summarize, our data indicate a close relationship between medial frontal theta, central beta activity, and the executive components of WM. Both oscillatory indices of executive processes, manifested in the theta (as we expected) and beta (unexpectedly) frequency bands, were related to behavioral performance in a verbal WM task. In contrast, we could not replicate the data indicating the important role of posterior alpha in WM performance. We can conclude that the ability to control attention plays a larger role in individual differences in WM than the capacity of the short-term storage.

## Methods

### Participants

186 individuals participated initially in the study. Eight participants with overall performance below 60% were excluded. Furthermore, a subsequent analysis revealed 22 EEG records with an excessive number of artefacts (less than 12 clean trials in any condition). Thus, 156 participants (82 females, mean age = 21.23, SD=3.22) were included in the final sample. The participants had normal or corrected-to-normal vision and self-reported no history of neurological or mental diseases. All the participants were Russian native speakers. The experimental protocol was approved by the Ural Federal University ethics committee, and conducted in accordance with the Declaration of Helsinki. Informed consent was obtained from all participants.

### Stimuli

Sets of Cyrillic alphabet letters written in capitals were used as stimuli. All letters (vowels and consonants) had been selected randomly from the alphabet, had random order, and no repetitions in the sets. The pool of stimuli comprised 1200 sets of letters (400 per level of load) which were manually checked to not form any meaningful words. An analogue using Latin letters and English words is shown in Figure 1.

A trial always began with an exclamation mark presented for 200 ms, which was followed by a fixation cross for 3000 ms. Participants were instructed to fixate on the cross whenever it appeared. Next, the word “forward” or “alphabetical”, presented for 600 ms, instructed the participants whether they would have to maintain in memory the original set as it was presented (retention task) or, first, mentally reorganize the letters into the alphabetical order and then maintain the result in memory (manipulation task). After that, sets of 5, 6 or 7 letters were demonstrated for 3000 ms followed by a delay period where a fixation cross was demonstrated for 6700 ms. At the end of the delay period, a randomly chosen letter from the previously presented set appeared on the screen together with a digit that represented the serial number of this letter. The letter-digit probe was presented for 1000 ms. Depending on the task, the participants indicated whether the probe was on the corresponding position either in the original set (retention task), or in the set resulted from alphabetical reordering (manipulation task). The participants were asked to press one of the computer mouse buttons (left or right) if the probe was correct and the other button otherwise. The two buttons were attributed to the correct and wrong probes in a counterbalanced order. The probe was correct in 50 % of the trials, and the order of correct and incorrect probes was randomized. The next trial started after a blank interval that varied between 5000 and 5500 ms.

The experiment entailed six different conditions: maintenance in memory of 5, 6 or 7 letters in the alphabetical (manipulation condition) or forward (retention condition) order. Each condition had 20 consecutive trials. These six blocks of 20 trials were presented in a random order. Two practice blocks with 3 and 6 trials respectively were given shortly before the main experiment.

During the experiment, the participants were seated in a comfortable armchair in front of a computer screen in a dark room. Stimuli were presented in white color on a black background in the center of the screen by using PsyTask software (Mitsar Ltd.). The distance to the screen was 1 m and the size of the letters was 1.2 × 1.2°.

### EEG recording and preprocessing

The EEG was recorded from 19 electrodes arranged according to the 10-20 system using Mitsar-EEG-202 amplifier with averaged earlobes reference. Two additional electrodes were used for horizontal and vertical EOG. EEG data were acquired with 500 Hz sampling rate and 150 Hz low-pass filter.

The procedure of EEG artifacts suppression and removal was conducted in two steps. At the first step, in order to suppress ocular activity artifacts, the independent component analysis (ICA) was performed using AMICA algorithm ^50^. The components clearly related to blinks and eye movements were identified and removed after visual exploration of the data. At the second step, epochs still containing artefacts were visually identified and discarded.

EEGLAB toolbox ^51^ for MATLAB was used for the data preprocessing.

### Time-frequency analysis

Before the time-frequency analysis, 1 Hz high-pass, 45 Hz low-pass and 50 Hz notch filters were applied with the EEGLAB firfilt function. Then, epochs in [-14200 2200 ms] interval where 0 is the onset of the probe were created.

The EEG time series in each epoch were convolved with a set of complex Morlet wavelets. The wavelets were defined as the product of a complex sine wave and a Gaussian window – 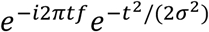, where t is time, f is frequency. σ is the width of the Gaussian, which set according to *n/(2πf)*, where *n* is the number of cycles – the parameter defining the time-frequency precision trade-off. The frequency *f* increased from 1 to 45 Hz in 45 linearly spaced steps, and the number of cycles *n* increased from 3 to 12 in 45 logarithmically spaced steps. From the resulting complex signal, the power of each frequency at each time point was obtained. The power was baseline-normalized by computing the percent change of the power in respect to [−11500 −10500] ms interval (i.e. “Baseline” in Figure 1).

The time-frequency analysis was performed by means of the Fieldtrip toolbox ^52^.

### Statistics

In order to decrease the number of factors employed in statistical calculations we defined frequency-channels regions of interest (ROI) with maximal representation of certain frequencies in certain group of channels. Thus, theta (4-8 Hz) had the maximal power at Fz, alpha (9-14 Hz) at posterior channels (left: T5, P3, O1; right: T6, P4, O2) and beta (16-22 Hz) at central channels (C3, Cz, C4) (see Supplementary materials for time-frequency maps at each channel).

Then spectral power was averaged in the ROIs in a 6 s time interval corresponding to the delay period (see Figure 2). We excluded the initial 700 ms from the onset of Delay period. As can be seen in Figure 2 and Supplementary Figure 1, the first few hundred milliseconds after the onset of the delay contain evoked response activity that could distort the frequency data if not rejected ^53–55^.

For statistical analysis of behavioral data, we employed repeated-measures analysis of variance (RM ANOVA) with factors Task (2 levels: Manipulation, Retention) and Load (3 levels: 5, 6, 7 letters to memorize).

For EEG data analyses we employed linear mixed-effects models (LME). As recommended by Barr et al. ^56^, first, we always tried to fit the maximal model. In the case of the convergence problem, we first tried to fit the model with main effect random slopes (no interactions). If it converged, next, we again followed the advice by Barr ^57^ and reduced the maximal random effect structure keeping the highest-order interaction slope (e.g., Task x Load x Hemisphere for alpha rhythm) and slopes that showed significant interactions with Performance at the previous step. Before fitting the models, continuous independent variables were centered around zero, and qualitative variables were effect coded.

For beta and theta rhythms the maximal model included Task and Load, Performance and all possible interactions as fixed effects, Participant as a grouping random intercept effect, and combination of Task, Load and their interactions as random slopes. For alpha activity fixed and random effects of Hemisphere (2 levels: left, right) were additionally included, because alpha frequently shows an asymmetry during delay period (Pavlov & Kotchoubey, 2020). Performance was calculated as the mean percentage of correct answers averaged over all conditions. In this case Performance plays the role of a personal trait and allows a straightforward interpretation of possible interactions with the other fixed effects. Single trial EEG data entered the analysis.

lme4 package ^58^ and function lmer in *R* was used to fit the models. For the calculation of p-values we used lmerTest package ^59^ with the default Satterthwaite’s degrees of freedom approximation.

We decided to use the alpha level of 0.005, as such a threshold exhibits higher evidential value and may also help to improve reproducibility of newly discovered effects (e.g. see Benjamin et al. ^60^ but also Miller & Ulrich ^61^ for another opinion).

Reliability estimates (Cronbach’s alpha) for behavioral and EEG variables in the manipulation and retention tasks are reported in Supplementary Table S4.

All statistical calculations were performed in *R*, version 3.6.3 ^62^.

## CRediT authorship contribution statement

YGP: Conceptualization, Funding acquisition, Investigation, Data curation, Formal analysis, Project administration, Resources, Supervision, Visualization, Methodology, Software, Writing - original draft, Writing - review & editing.

BK: Methodology, Writing - review & editing.

## Potential Conflicts of Interest

Nothing to report.

### Acknowledgements

The study was supported by Russian Foundation for Basic Research (RFBR) #19-013-00027. We acknowledge support by the Deutsche Forschungsgemeinschaft and Open Access Publishing Fund of University of Tübingen.

## Data availability

The datasets generated during and/or analyzed during the current study are available from the corresponding author on request, without undue reservation.

