## Supplementary for "The electrophysiological underpinnings of variation in verbal working memory capacity"

### Supplementary Methods

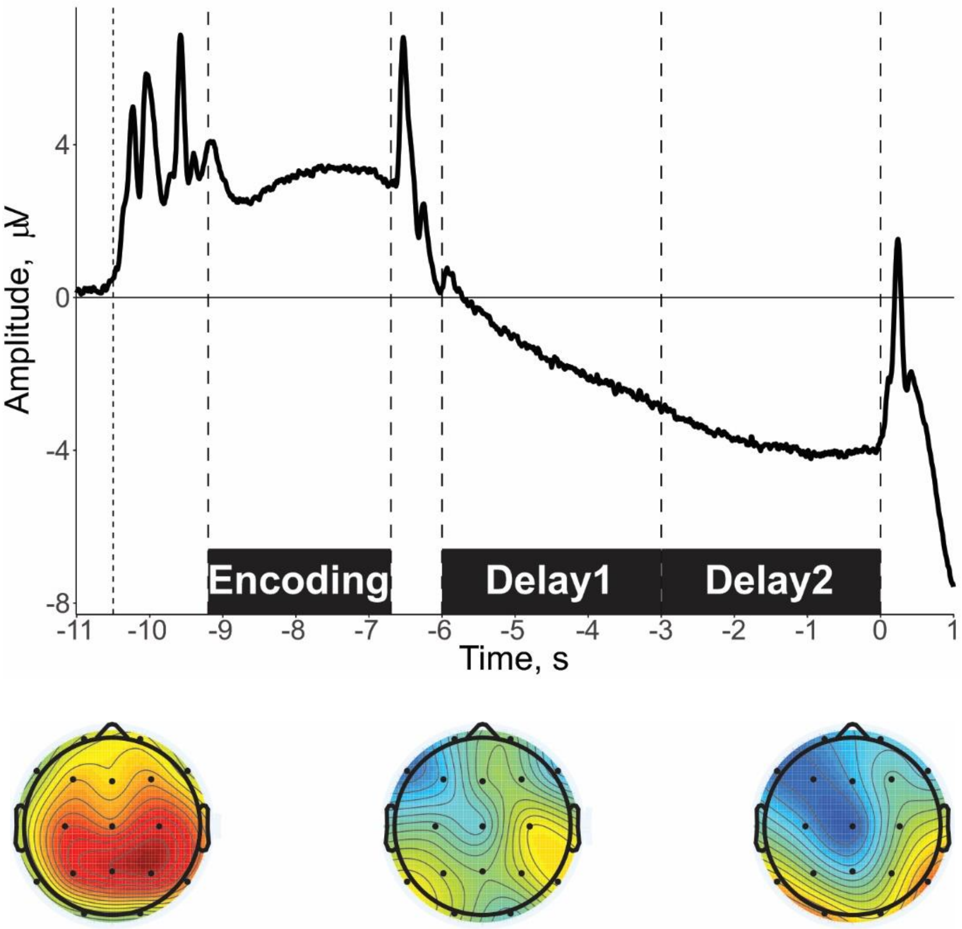


Figure S1 – Grand average waveforms of event-related potentials (ERPs) over all conditions in Cz channel. The figure shows two spikes in the period of 700 ms at the beginning of the delay that was excluded from the analysis. Topographical plots in the bottom panel represent grand average ERPs in Encoding [last 2.5 s of the encoding period], Delay1 [first 3 s of Delay], and Delay2 [second 3 s of Delay], respectively. The topoplots were created in MATLAB R2016b (MathWorks Inc.). The waveforms were visualized with ggplot2 package (Wickham, 2016) in R v.3.6.3.

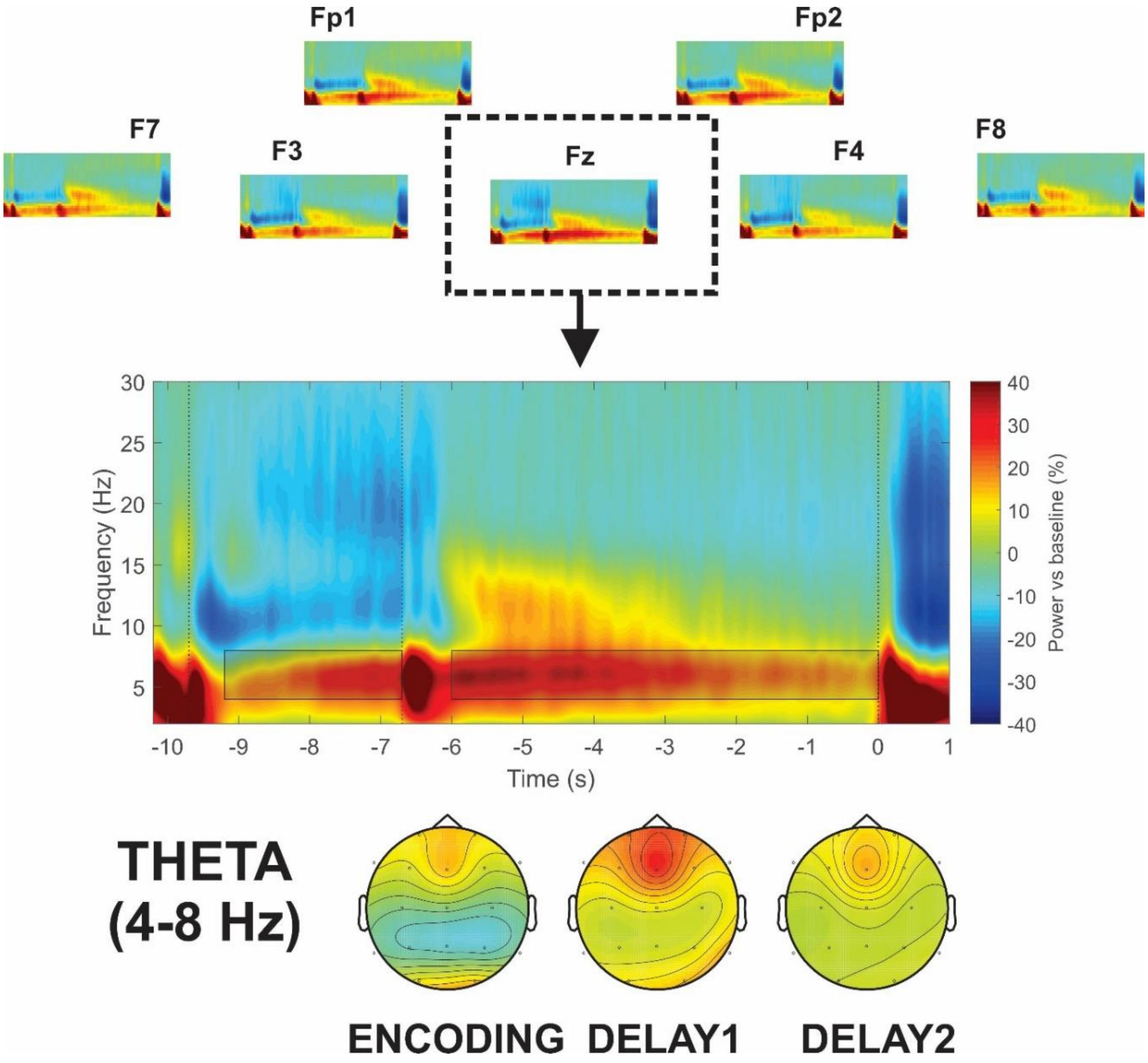


Figure S2 – Top panel: Time-frequency maps in individual channels. The power scale is the same as in the middle panel and bottom panels. Middle panel: Fz theta grand average. Two boxes mark the time windows: Encoding and Delay. The encoding period is shown to demonstrate how the time-frequency windows were determined. Bottom panel: Topographical maps of averaged baseline-normalized theta (4-8 Hz) power in Encoding [last 2.5 s of the encoding period], Delay1 [first 3 s of Delay], and Delay2 [second 3 s of Delay], respectively. Two boxes mark the time windows: Encoding and Delay. The time-frequency maps and topoplots were created in MATLAB R2016b (MathWorks Inc.)

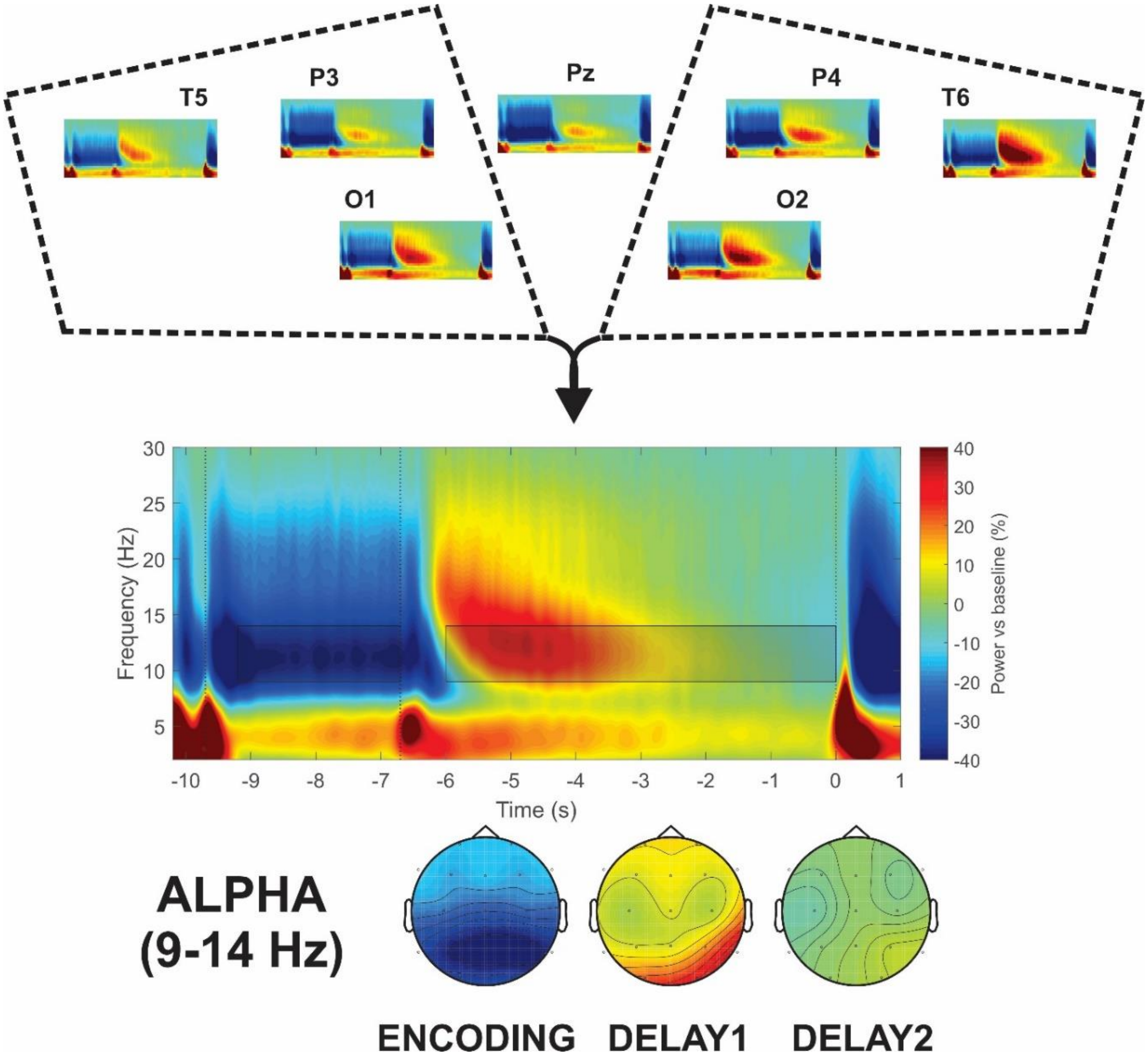


Figure S3 – Top panel: Time-frequency maps in individual channels. The power scale is the same as in the middle and bottom panels. Middle panel: Alpha ROI (Left: T5, P3, O1; Right: P4, T6, O2) grand average. Two boxes mark the time windows: Encoding and Delay. The encoding period is shown to demonstrate how the time-frequency windows were determined. Bottom panel: Topographical maps of averaged baseline-normalized alpha (9-14 Hz) power in Encoding [last 2.5 s of the encoding period], Delay1 [first 3 s of Delay], and Delay2 [second 3 s of Delay], respectively. The time-frequency maps and topoplots were created in MATLAB R2016b (MathWorks Inc.)

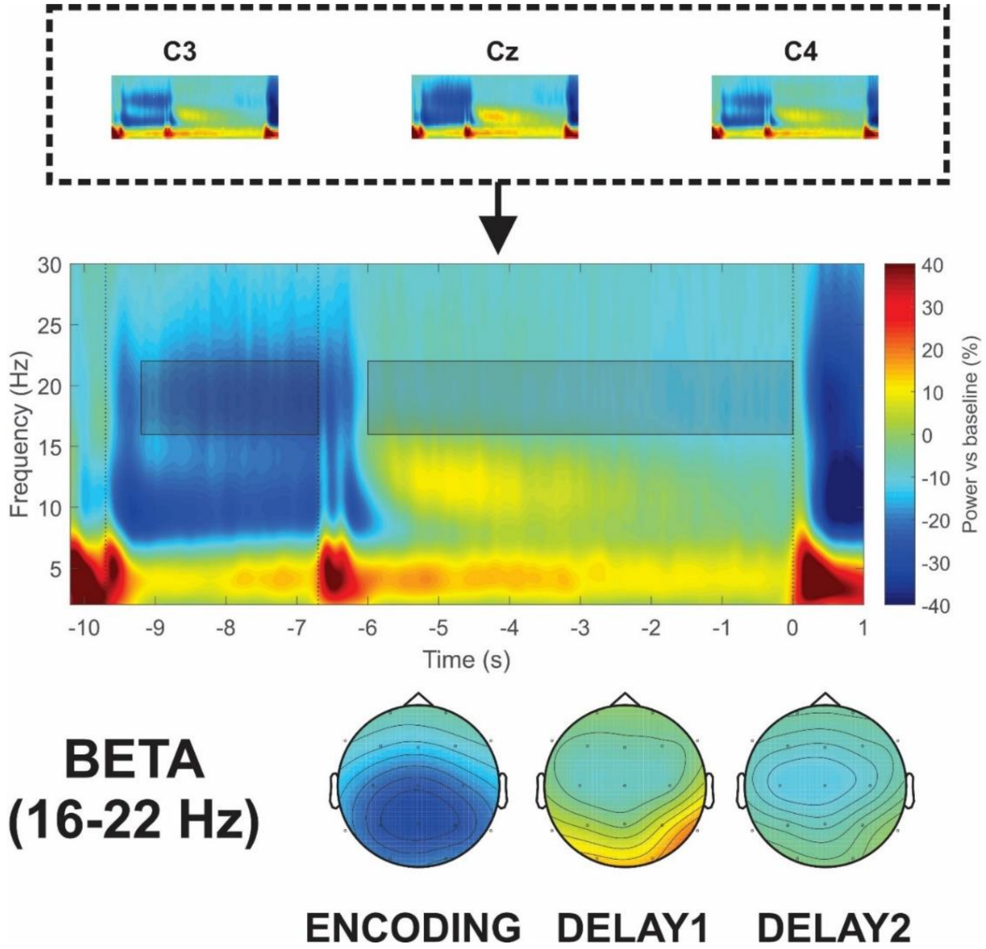


Figure S4 – Top panel: Time-frequency maps in individual channels. The power scale is the same as in the middle panel and bottom panels. Middle panel: Beta ROI (C3, Cz, C4) grand average. Two boxes mark the time windows: Encoding and Delay. The encoding period is shown to demonstrate how the time-frequency windows were determined. Bottom panel: Topographical maps of averaged baseline-normalized beta (16-22 Hz) power in Encoding [last 2.5 s of the encoding period], Delay1 [first 3 s of Delay], and Delay2 [second 3 s of Delay], respectively. The time-frequency maps and topoplots were created in MATLAB R2016b (MathWorks Inc.)

### Supplementary results

#### Linear mixed-effect models

The models formulas in the results section are written using the syntax of lme4 package to make them more comprehensible. For example, the formula for theta spectral power effects related to WM performance will be presented as follows:

Power ~ Task * Load * Performance + (Load*Task |Participant)

Power is the dependent variable. Task, Load, Performance before the parentheses are fixed effects; Load, Task in the parentheses on the left from | sign are random slopes; and Participant is random intercept, i.e., the grouping variable.

The maximal model for theta successfully converged. The model's description:

*Power ~ Task * Load * Performance + (Load*Task |Participant)*

See Table S1 for LMM of theta statistical output.

Table S1 - Theta statistics in LME

| | *β* | *SE* | *t* | *p* |
|---|---|---|---|---|
| (Intercept) | 22.78 | 2.61 | 8.72 | < 0.001 |
| Task | -3.41 | 1.05 | -3.25 | 0.001 |
| Load | -1.37 | 0.8 | -1.72 | 0.088 |
| Performance | -1.1 | 2.61 | -0.42 | 0.673 |
| Task:Load | 4.36 | 1.79 | 2.43 | 0.016 |
| Task:Performance | -3.07 | 1.05 | -2.93 | 0.004 |
| Load:Performance | 2.04 | 0.8 | 2.55 | 0.012 |
| Task:Load:Performance | -2.27 | 1.79 | -1.26 | 0.208 |

The final model for alpha that converged successfully:

*Power ~ Hemisphere * Task * Load * Performance + (Hemisphere+ Load+Task+Task:Load+Task:Load:Hemisphere |Participant)*

As shown in Table S2 no significant main effects or interactions with Performance were found in the alpha frequency band.

Table S2 – Alpha LME

| | *β* | *SE* | *t* | *p* |
|---|---|---|---|---|
| (Intercept) | 10.96 | 2.12 | 5.16 | < 0.001 |
| Hemisphere | 5.19 | 0.65 | 8.04 | < 0.001 |

| | | | | |
|---|---|---|---|---|
| Task | 2.75 | 0.95 | 2.9 | 0.004 |
| Load | 1.58 | 0.9 | 1.74 | 0.083 |
| Performance | -1.81 | 2.12 | -0.85 | 0.395 |
| Hemisphere:Task | -1.03 | 0.27 | -3.79 | < 0.001 |
| Hemisphere:Load | 0.27 | 0.33 | 0.8 | 0.423 |
| Task:Load | -1.3 | 1.05 | -1.24 | 0.216 |
| Hemisphere:Performance | 1.18 | 0.65 | 1.83 | 0.069 |
| Task:Performance | 0.71 | 0.95 | 0.75 | 0.454 |
| Load:Performance | 0.61 | 0.9 | 0.68 | 0.499 |
| Hemisphere:Task:Load | 0.21 | 0.38 | 0.55 | 0.583 |
| Hemisphere:Task:Performance | -0.08 | 0.27 | -0.31 | 0.756 |
| Hemisphere:Load:Performance | -0.12 | 0.33 | -0.35 | 0.723 |
| Task:Load:Performance | 1.84 | 1.05 | 1.76 | 0.08 |
| Hemisphere:Task:Load:Performance | 0.34 | 0.38 | 0.9 | 0.371 |

The maximal model for beta successfully converged. In *lmer* syntax the final model can be described by the following formula:

*Power ~ Task * Load * Performance + (Load*Task |Participant)*

Table S3 – Beta LME

| | *β* | *SE* | *t* | *p* |
|---|---|---|---|---|
| (Intercept) | -8.95 | 0.78 | -11.44 | < 0.001 |
| Task | 1.84 | 0.44 | 4.22 | < 0.001 |
| Load | 0.11 | 0.46 | 0.23 | 0.816 |
| Performance | -1.22 | 0.78 | -1.56 | 0.121 |
| Task:Load | -1.46 | 0.46 | -3.17 | 0.002 |
| Task:Performance | 1.51 | 0.44 | 3.47 | < 0.001 |
| Load:Performance | -0.76 | 0.46 | -1.64 | 0.103 |
| Task:Load:Performance | 0.44 | 0.46 | 0.97 | 0.336 |

### Correlation analysis

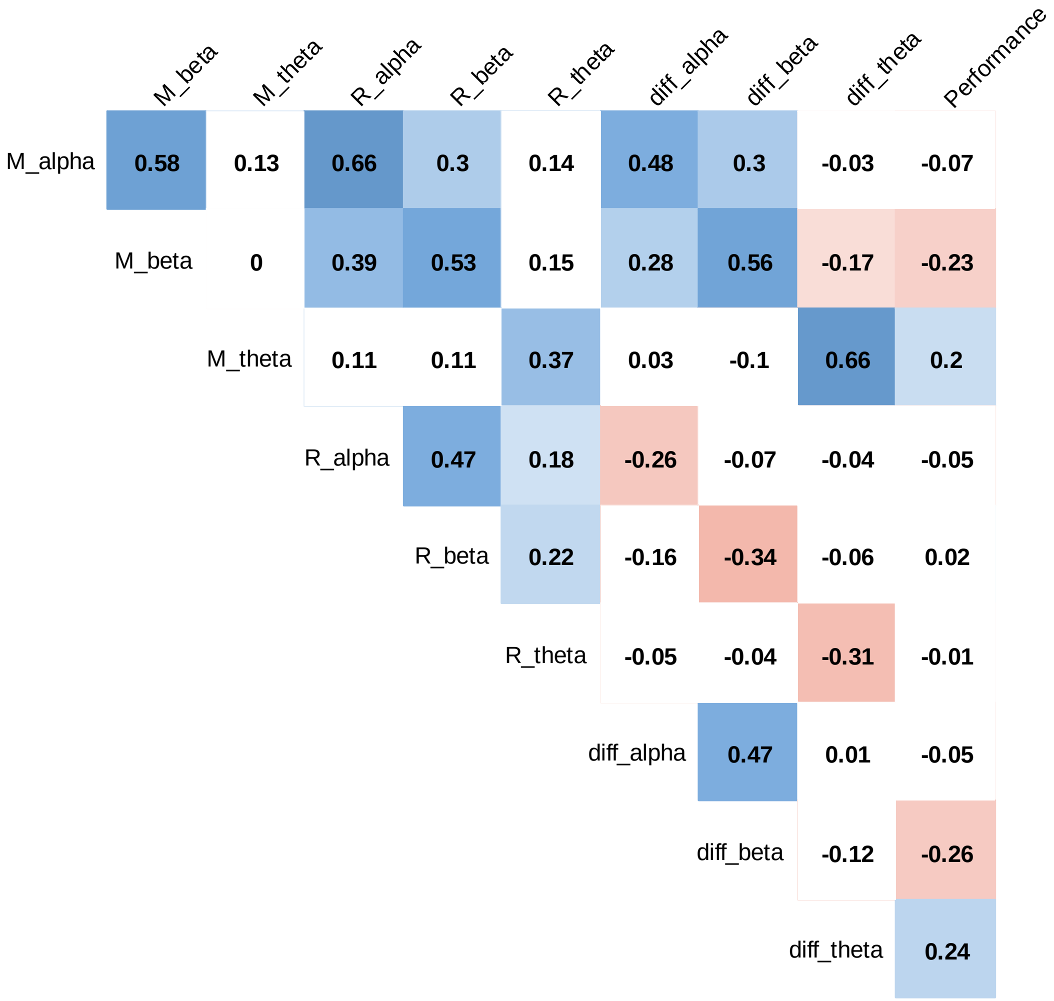


Figure S5 – Correlation matrix relating EEG indices and overall accuracy. The highlighted with color cells are significant correlations with p < .05. M – manipulation task, R – retention task, diff – difference score M-R, Performance – average accuracy in all conditions. The figure was created in corrplot package (Wei, & Simko, 2017) for R v.3.6.3.

### Reliability analysis

We used adjusted Cronbach's alpha for the difference score (Furr and Bacharach, 2013, formula below) and Cronbach's alpha for other measures.

$$R_d = \frac{S_x^2 R_{xx} + S_y^2 R_{yy} - 2r_{xy}S_xS_y}{S_x^2 + S_y^2 - 2r_{xy}S_xS_y},$$

where $s_x^2$ squared and $s_y^2$ are the variance of subject average scores, $s_x$ and $s_y$ are the standard deviation of subject averaged scores, $R_{xx}$ and $R_{yy}$ are the split-half internal consistency, and $r_{xy}$ is the correlation between the measures. The split-half internal consistency ($R_{xx}$ and $R_{yy}$ in the formula) was calculated using splitHalf function as implemented in *psych* package for R (Revelle, 2019).

Table S4 – Internal consistency estimates

| | | Cronbach's α |
|---|---|---|
| Retention | alpha | 0.948 |
| | beta | 0.938 |
| | theta | 0.94 |
| | accuracy | 0.7 |
| Manipulation | alpha | 0.965 |
| | beta | 0.952 |
| | theta | 0.956 |
| | accuracy | 0.679 |
| Manipulation - Retention difference score | alpha | 0.874 |
| | beta | 0.891 |
| | theta | 0.832 |
| | accuracy | 0.511 |
| Overall | alpha | 0.973 |
| | beta | 0.964 |
| | theta | 0.964 |
| | accuracy | 0.772 |

*Notes:* accuracy – percent of correct responses in a particular task; overall – consistency estimates for the measures averaged over all conditions; alpha, beta, theta – power in the frequency bands of interest (see Methods).
